# Evidence for glutamine synthetase function in mouse spinal cord oligodendrocytes

**DOI:** 10.1101/2021.03.29.437580

**Authors:** Lucile Ben Haim, Lucas Schirmer, Amel Zulji, Khalida Sabeur, Brice Tiret, Sandra Chang, Wouter H. Lamers, Myriam M. Chaumeil, David H. Rowitch

**Author notes:** Correspondence: Lucile Ben Haim, David H. Rowitch.

## Abstract

Glutamine synthetase (GS) is a key enzyme that metabolizes glutamate into glutamine. While GS is expressed by astrocytes of the central nervous system (CNS), expression in other glial lineages has been noted. Using a combination of reporter mice and cell typespecific markers, we show that GS is expressed in myelinating oligodendrocytes (OL) but not oligodendrocyte progenitor cells (OPC) of the mouse spinal cord abutting ventral horn motor neurons. To investigate the role of GS in mature OL, we used a conditional knockout (*cKO*) approach to selectively delete GS-encoding gene (*Glul*) in OL, which caused a significant decrease in glutamine levels on spinal cord extracts. We evaluated the effect on ventral spinal cord sensorimotor circuits and observed that *GS cKO* mice (*CNP-cre*^+^: *Glul^fl/fl^*) showed no differences in motor neuron numbers, size or axon density; OL differentiation and myelination in the ventral spinal cord at 1- and 6-months of age was normal. Interestingly, *GS cKO* mice showed an early and specific decrease in peak force while motor function remained otherwise unaffected. These findings provide evidence OL-encoded GS functions in spinal cord sensorimotor circuit.

**Main points:**

- Glutamine synthetase (GS) is highly expressed in oligodendrocytes (OL) of the mouse spinal cord.
- OL-specific GS loss of function causes transient decrease in peak force, but other substantial neurological dysfunction was not detected.

## Introduction

Glutamine synthetase (GS) is an ATP-dependent enzyme, which catalyzes the transformation of glutamate (Glu) and ammonium into glutamine (Gln) (Mates, Campos-Sandoval, Santos-Jimenez, & Marquez, 2019). It is involved in maintaining Gln physiological levels and in the rapid recycling of Glu through the “Glu-Gln cycle”, which is crucial for excitatory neurotransmission (Daikhin & Yudkoff, 2000; Norenberg & Martinez-Hernandez, 1979). In the central nervous system (CNS), GS has generally been considered an astrocyte-specific enzyme, especially in forebrain regions (Liang, Carlson, & Coulter, 2006; Papageorgiou et al., 2018). Loss-of- function of GS-encoding gene, *Glul*, leads to major cerebral abnormalities in humans, decreased Gln levels (Haberle et al., 2005) and early neonatal death in mice (He et al., 2010). Astrocyte-specific expression is altered in many pathological conditions including neurodegenerative diseases, hypoxia or epilepsy both at the gene and protein levels (Jayakumar & Norenberg, 2016; Rose, Verkhratsky, & Parpura, 2013; Xin et al., 2019). In addition to astrocytes, GS expression has also been reported in oligodendrocytes (OL) in the brain and in the spinal cord white matter (Bernstein et al., 2014; Cammer, 1990; Takasaki et al., 2010) but remains controversial (Anlauf & Derouiche, 2013). Given the key role of GS function in neuron-glia crosstalk, we investigated its precise expression and function in OL lineage cells in the mouse sensorimotor circuit, focusing on the spinal cord.

## Material and methods

### Mice

All mouse strains were maintained at the University of California, San Francisco (UCSF) pathogen-free animal facility and all animal protocols were approved by and in accordance with the guidelines established by the Institutional Animal Care and Use Committee and Laboratory Animal Resource Center under protocol number AN110094. *Aldh1l1-eGFP* (OFC789Gsat/ 011015-UCD) transgenic mice were generated by the GENSAT project (Gong et al., 2003) and *MOBP-eGFP* mice (IN1Gsat) were obtained from MMRRC and maintained as homozygous on a C57BL/6J background. *Glul^fl/fl^* mice were obtained from Dr. W. Lamers (Amsterdam University Medical Centers) (He et al., 2010). *CNP-cre* transgenic mice (Lappe-Siefke et al., 2003) were commercially available (Jackson Lab). *NG2-DsRed* mice (Tg(*Cspg4-DsRed.T1*)1Akik) were obtained from Dr. S. Fancy (UCSF). *Glul^fl/fl^* mice were bred with *CNP-cre: Glul^fl/+^. CNP-cre^-^: Glul^fl/fl^* or ^*fl*/+^ where used as controls and *CNP-cre^+^: Glul^fl/fl^* as *cKO*. For clarity, *CNP-cre^+^: Glul^fl/fl^* will be abbreviated as *GS cKO* in the manuscript. All mice were maintained on a 12 hr light/dark cycle with food and water available *ad libitum*. For all experiments, we used both male and female mice. The number of males and females in each analysis group was balanced as much as possible. No significant effect of sex was observed in data analyses.

### Human tissue

Human spinal cord samples were obtained through the UK Multiple Sclerosis Tissue Bank at Imperial College, London. Three snap-frozen control human tissues from deceased individuals without spinal cord pathology (**Supp. Table 1**) were obtained via a prospective donor scheme following ethical approval by the National Research Ethics Committee in the UK (08/MRE09/31). We have complied with all relevant ethical regulations regarding the use of human postmortem tissue samples.

### Immunofluorescence stainings

Mice were transcardially perfused with 4% paraformaldehyde. After post-fixation, samples were cryoprotected in 30% sucrose for 48h at 4°C and embedded in optimal cutting temperature compound (Tissue-Tek). Spinal cords were cut using a cryostat (CM3050S, Leica) and either collected on superfrost slides (VWR) (14-16μm-thick sections) or in a P24 well plate, in cryoprotecting solution (30μm-thick sections). Free-floating sections were stored in cryoprotecting solution at −20°C. Just before processing, sections were rinsed twice during 48h in 0.1M phosphate buffer (PB) to remove storing solution. Sections were subjected to antigen retrieval in 1X Dako Target solution for 2 min at 95°C, permeabilized and blocked in 0.1M phosphate buffered saline (PBS)/0.2% Triton X (TX)-100/10% horse serum (HS) for 1h at room temperature (RT). Primary antibody incubations were carried out overnight□at 4□°C. After washing in 0.1M PBS, sections were incubated with secondary antibodies diluted in 0.1M PBS/ 0.2% TX-100/10% HS for 1h, RT. Goat or donkey Alexa fluochrome-tagged secondary IgG antibodies were used for primary antibody detection. Slides were mounted with DAPI Fluoromount-G (SouthernBiotech).

For human tissue analysis, frozen sections were fixed 5 min in 100% ice-cold methanol, rinsed in 0.1M PBS/0.1% TX-100/5% HS and incubated overnight at 4°C with primary antibodies in 0.1M PBS/0.1% TX-100/5% HS. Sections were then rinsed and incubated with the secondary antibodies in 0.1M PBS/0.1% TX-100/5% HS for 30 min. Sections were then rinsed and mounted using DAPI Fluoromount-G. Primary antibodies used included: goat anti-ChAT (AB144P, Millipore, 1:200), rat anti-GFAP (13-0300, Invitrogen, 1:1000), rabbit anti-GS (G2781, Sigma, 1:1000, antibody directed against C-term AA 357-363 with N-terminally added Lys), mouse anti-GS (MAB302, clone GS-6, Millipore, 1:1000), mouse anti-NeuN (MAB377, Millipore, 1:1000), guinea pig anti-VGLUT1 (AB5905, Millipore, 1:5000), guinea pig anti-VGLUT2 (AB2251, Millipore, 1:5000), goat anti-Olig2 (AF2418, R&D Systems, 1:50), mouse anti-APC (clone CC1, OP80, 1:300, Millipore Sigma), mouse anti-NOGO-A (clone 11C7, gift from M.E. Schwab, 1:3,000), rat anti-MBP (ab7349, Abcam, 1:500), mouse anti-Neurofilament H (NF-H), phosphorylated neurofilament (clone SMI312, 837904, Biolegend, 1:1,000), rabbit anti-IBA1 (019–19741, Wako, 1:500).

### Image acquisition and analysis

Images were acquired on a Zeiss Axio Imager (10x objective), a Leica TCS SPE or SP8 confocal microscopes (20x or 40x objectives). All confocal pictures are z-stack confocal images. The mean fluorescence intensity (immunoreactivity) was measured on images from samples processed in a strictly identical manner and acquired with the same parameters (laser power and gain). Image analysis was performed using ImageJ on 10x epifluorescence or 40x confocal images. For epifluorescence images, cell numbers, mean or maximum fluorescence intensity were quantified on an average of 6-8 serial spinal cord hemisections. For confocal images, quantifications were performed on 3-4 fields in the ventral horn, on 3 serial sections. MN soma size was measured as described previously (Kelley et al., 2018). The % of immune-positive area for GFAP and IBA1 was measured using the threshold function on Image J on 8-bit stacked confocal images.

### Quantitative RT-PCR

RNA was isolated using Trizol reagent (Invitrogen), DNase-digested to remove genomic DNA and purified using the RNAeasy Kit (Qiagen) according to manufacturer’s instructions. Complementary DNA was generated using Superscript III (Invitrogen) and random hexamers. qPCR was performed by using SYBR Green master mix (Roche) in Light-cycler 480 (Roche) with specific primers designed for amplicons of 75-150 bp using Primer 3. The following primer sequences were used: *Glul* (F: GGGCTACTTTGAAGACCGTC; R: TTCGTCGCCTGTTTCGT), *Pdgfrα (F: GGCCAGAGACATCATGCACGATTC; R: TCAGCGTGGTGTAGAGGTTGTCGAA*), *Mbp* (F: CCCAAGGCACAGAGACACGGG; R: TACCTTGCCAGAGCCCCGCTT), and *18S* (F: GTTCCGACCATAAACGATGCC; R: TGGTGGTGCCCTTCCGTCAAT). *18S* was used as housekeeping gene and melting curve analyzed to ensure correct and specific amplification.

### Western blot

Mouse brainstem tissue (pons and medulla oblongata) samples were dissected. Sample lysis was performed in RIPA buffer (Thermofisher) in the presence of protease and phosphatase inhibitors (Cell signaling). Samples concentration was determined with the Bradford method and protein migration and gel transfer was performed as described previously (Kelley et al., 2018). After blocking in Odyssey^®^ Blocking Buffer (PBS) (Li-Cor) for 1h, RT, primary antibodies were incubated overnight at 4°C onto the western blot membrane. The following antibodies were used: rabbit anti-GS, rat anti-MBP and mouse anti-III *β*-Tubulin (MMS435P, Covance, 1:10000). IRDye^®^ Goat anti-mouse and anti-rabbit (680 and 800) fluorescent secondary antibodies (Li-cor) were used for protein detection on the Odyssey Cxl imaging system.

### Ex vivo ^1^H NMR

Metabolites from *GS OL cKO* and control 2 month-old mice whole spinal cord were extracted using equal parts methanol-water-chloroform, as previously described (Tyagi, Azrad, Degani, & Salomon, 1996). Briefly, animals were euthanized and 89 ± 5mg of spinal cord tissue collected and snap-frozen in liquid nitrogen. Using a mortar and pestle, frozen tissue was homogenized in −20°C methanol. Equal parts −20°C chloroform and 4°C H2O were homogenously mixed, and the fractions separated by centrifuging at 125g at 4°C. The methanol fraction was collected, 11.7mM Trimethylsilylpropanoic acid (Acros Organics) added, and the mixture lyophilized. The resultant extracts were reconstituted in 420μL distilled H2O and samples were scanned on an 500Mhz NMR system (Bruker) with a 1D pulse acquire zgpr sequence sequence (NS = 16, DS 2, 32k datapoint, TR 2sec). N-acetyl aspartate (NAA), glutamine and glutamate were fitted and quantified using Chenomx NMR Suite (Chenomx Inc) with reference to the Human Metabolomics Database (Wishart et al., 2018).

### Behavioral analysis

All behavioral experiments were performed at the UCSF Neurobehavioral Core for Rehabilitation Research (UCSF). Mice were habituated to the experimenter through handling, weighting and cage change every week prior to behavioral testing. Experiments were all performed at the same time of the day for each day of testing. The rotarod test (Ugo Basile) was performed to assess motor coordination and motor learning. Mice were placed onto a rotating rod accelerating from 0-40 rotations per minute during 5 minutes. Animals were tested 3 times per day for 3 consecutive days and the latency to fall from the apparatus at each trial was measured. Forelimb grip strength was measured using a grip strength apparatus (Bioseb). The grip strength of each mouse was measured on 3 trials per day for 3 consecutive days and the average peak force per animal was computed. The open field test (Kinder-Scientific) was used to assay overall locomotor activity. Mice were placed to freely explore an open arena connected to a video tracking software for 10 min. Data collected includes distance traveled and number of rearings.

### OPC immunopanning

Cells were isolated from neonatal mouse cortices using anti- PDGFRα (anti-CD140a, 558774, BD Biosciences) immunopanning antibody for positive selection of OPC (Yuen et al., 2014). Briefly, OPC were immunopanned from P7-P9 mouse cortices and plated on poly-D-lysine coverslips (Neuvitro). Cells were kept in proliferation media (PDGF-AA, CNTF, and NT3; Peprotech) at 10% CO_2_ and 37°C. After two days in proliferation media, differentiation was induced by changing media to contain CNTF and triiodothyronine (T3; Sigma). Mycoplasma contamination testing was negative.

### Bioinformatic analysis of existing data setss

Normalized and scaled data setss (feature-barcode matrices) from published mouse spinal cord single-cell RNA sequencing (scRNA-seq) (Blum et al., 2021) and human leukocortical single-nucleus RNA-sequencing (snRNA-seq) (Schirmer et al., 2019) and tissues were obtained (https://cells.ucsc.edu/ and http://spinalcordatlas.org/) and used without any processing. In addition, another mouse CNS scRNA-seq data sets (Marques et al., 2016) was reconstructed using the available count matrix and corresponding metadata (GSE75330). It should be noted that 16 cells in the metadata and matrix did not match, so these cells were omitted from the downstream analysis (**Supp. Table 2**). Briefly, the count matrix was transformed using SCTransoform (Hafemeister & Satija, 2019) as part of the Seurat toolkit (Stuart et al., 2019). The dimensionality of the data was reduced using principal component analysis (PCA) and the first 18 PCs were used for downstream clustering analysis. Cells were annotated using inferred cell identities provided by the authors in the metadata.

### Statistical analysis

We pre-determined group sample size based on previously published studies. Individual experimental values are shown on bar plots and no data were excluded from analysis. All bar graphs are expressed as mean ± SEM. In box-and-whisker plots, center lines indicate medians, box edges represent the interquartile range, and whiskers extend down to the 10th and up to the 90th percentiles of the distribution. Data sets normality was assessed using the Shapiro-Wilk test. Two-tailed unpaired parametric or non-parametric (Mann-Whitney test) t-tests were performed for two group comparisons. Data sets with n = 3 were analyzed with parametric tests as the non-parametric equivalents rely on ranking, which is not reliable for small sample size (GraphPad Prism 8). Repeated measures (genotype*time) ANOVA test or mixed effects model (if missing values) were to analyze grip strength and rotarod experiments. Level of significance was determined as described in the individual figure legends. P values were designated as follows: *p<0.05, **p<0.01, ***p<0.001. All statistical tests were performed using GraphPad Prism 9 (GraphPad Software).

## Results

### Macroglial-specific GS expression in the mouse spinal cord

To establish the precise pattern of GS expression in macroglial cells (astrocytes and OL lineage cells), we first used immunofluorescence detection in transgenic reporter mice for both astrocytes (*Aldh1L1- eGFP* mice) and mature myelinating OL (*MOBP-eGFP* mice) (**Figure 1A-D**). We indeed found that almost all *Aldh1L1-eGFP^+^* cells were also GS^+^ with a predominant expression in thin cytoplasmic processes (**Figure 1A, B**). In addition, almost 80% of *MOBP-eGFP^+^* OL in the ventral grey matter also expressed GS in their cell bodies, which accounting for ~50% of all GS^*+*^ cells (astrocytes + OL) (**Figure 1C, D**). We further confirmed that GS was expressed in mature OL identified by co-expression of CC1 and Olig2 (**Figure 1E-G**). This expression pattern was observed using two antibodies from different suppliers (**Supp. Figure 1**). The onset of GS expression in CC1^+^ OL was between postnatal day (P) 14 and 40, as it was only expressed in astrocytes at P14 (**Figure 1H, I**). These findings indicate that GS is expressed prominently in astrocytes and mature OL in mouse spinal cord.

**Figure 1.**
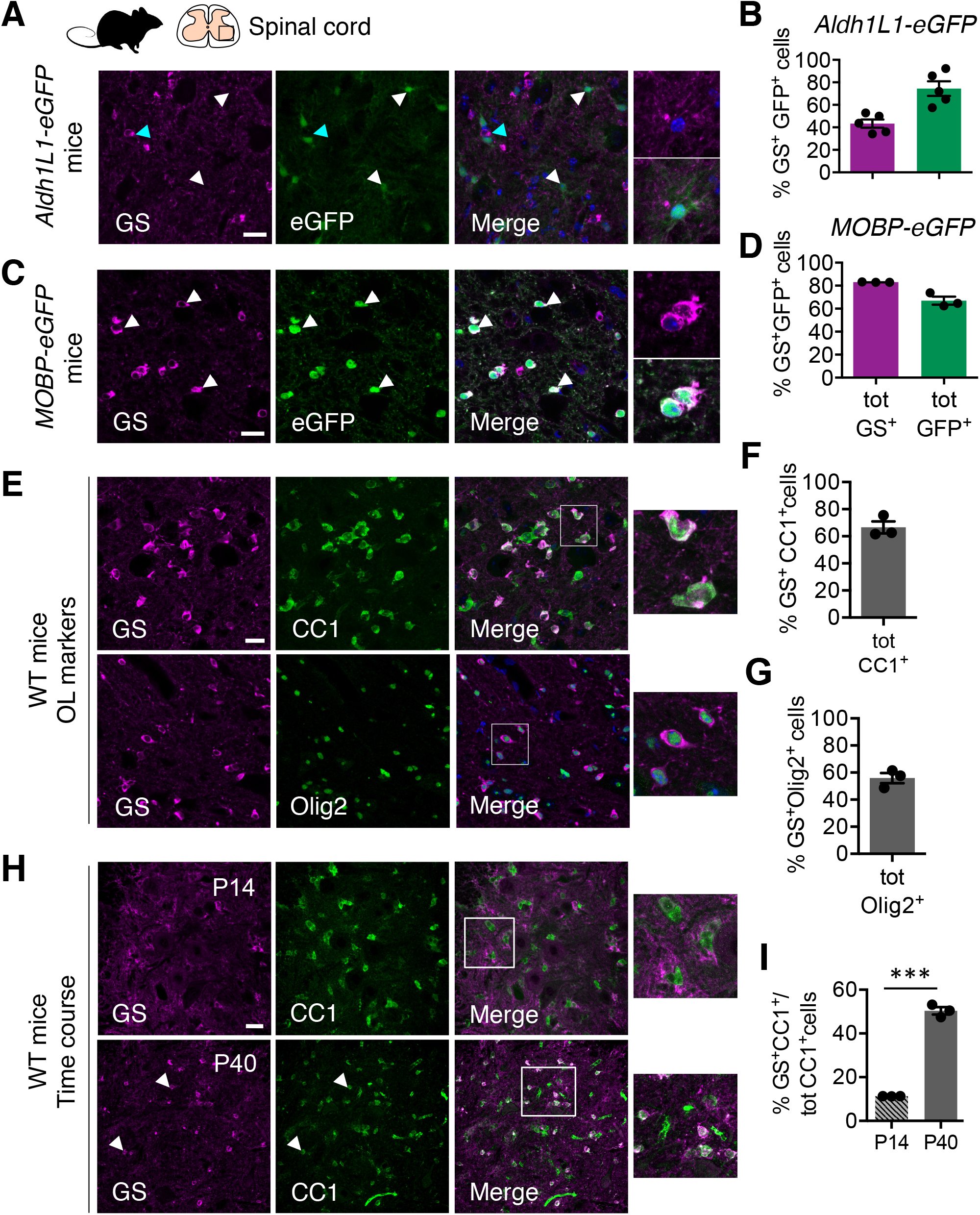
GS is expressed in astrocytes and mature OL in the mouse spinal cord. **A**, Confocal images of spinal cord sections from astrocyte reporter *Aldh1L1-eGFP* mice immunostained with GS (magenta). Note that GS expression in astrocytes is mainly located in thin cytoplasmic processes as compared to the soma, which are weakly GS^+^ (white arrowheads and high magnification). Cells with high somatic GS^+^ signal do not express *Aldh1L1-eGFP* (cyan arrowheads). **B**, Quantification of the % of GS^+^ astrocytes relative to total GS^+^ cells and total *Aldh1L1-eGFP* astrocytes. **C**, Confocal images of spinal cord sections from OL reporter *MOBP-eGFP* mice immunostained with GS (magenta) showing that a majority of *MOBP-eGFP* OL cell bodies express GS (white arrowheads and high magnification). **D**, Quantification of the % of GS^+^ OL in the total GS^+^ cell population (astrocytes and OL) and the total *MOBP-eGFP^+^* OL cell population. **E**, Confocal images of spinal cord sections of adult WT mice co-stained with GS (magenta) and OL markers CC1 (green) (top) and Olig2 (green) (bottom) showing that GS is expressed in CC1^+^ and Olig2^+^ OL. **F, G**, Quantification of the % GS^+^CC1^+^/total CC1^+^ (**F**) and GS^+^Olig2^+^/total Olig2^+^ cells (**G**). **H**, Confocal images of spinal cord sections from WT mice co-stained with GS (magenta) and CC1 (green) at postnatal age (P)14 (top) and P40 (bottom). GS^+^CC1^+^ cells (white arrowheads) are only detected in P40 but not P14 samples. **I**, Quantification of the % of GS^+^CC1^+^/ tot CC1^+^ OL at P14 and P40. Data are expressed as mean ± SEM (n = 3 - 4 mice). Unpaired t-test (t = 23.40, df = 4), ***p < 0.001. Scale bar: 20μm.

We next investigated whether GS was also expressed in OPC. We performed immunofluorescence co-stainings of GS with the OPC marker PDGFRα on WT mice or on *NG2-DsRed* OPC reporter mice spinal cord sections. We found no expression of GS in OPC (**Figure 2A**). Furthermore, using OPC immunopanning, we found that *Glul* transcripts levels of were higher in differentiation (mature OL) as compared to proliferation (OPC) conditions. *Pdgfrα* and *Mbp* transcripts levels were used as positive control markers for proliferation (OPC) and differentiation (OL) conditions, respectively (**Figure 2B**). We analyzed *Glul* expression in cells of the OL lineage from a published single cell RNA sequencing data sets (scRNA-seq; mouse CNS; Marques et al. (2016)) (**Figure 2C)**. In keeping with our results at the transcript and protein levels, *Glul* was enriched in mature OL (MOL) clusters 1 to 6 and its expression profile clearly resembles the mature myelinating OL marker, *Mobp* (**Figure 2D)**. By constrast, *Glul* expression level was low in newly formed OL and OPC clusters, identified with *Pdgfra* (**Figure 2D)**. Furthermore, we found no expression of GS in NeuN^+^ neurons or in IBA1^+^ microglia at the protein level (**Figure 2D, E**). These findings were consistent with *Glul* enrichment analysis in a published single nuclei (sn) RNA-seq data set from adult mouse spinal cord (Blum et al., 2021) (**Figure 2F**). We found that *Glul* was expressed in two main cell populations: mature OL, identified with *Mobp* and astrocytes (AS), identified with *Aqp4* expression. Some endothelial cells (*Flt1*) express *Glul* transcripts but it was not detected in microglia (*Cx3Cr1*) and neurons (*Rbfox3*).

**Figure 2.**
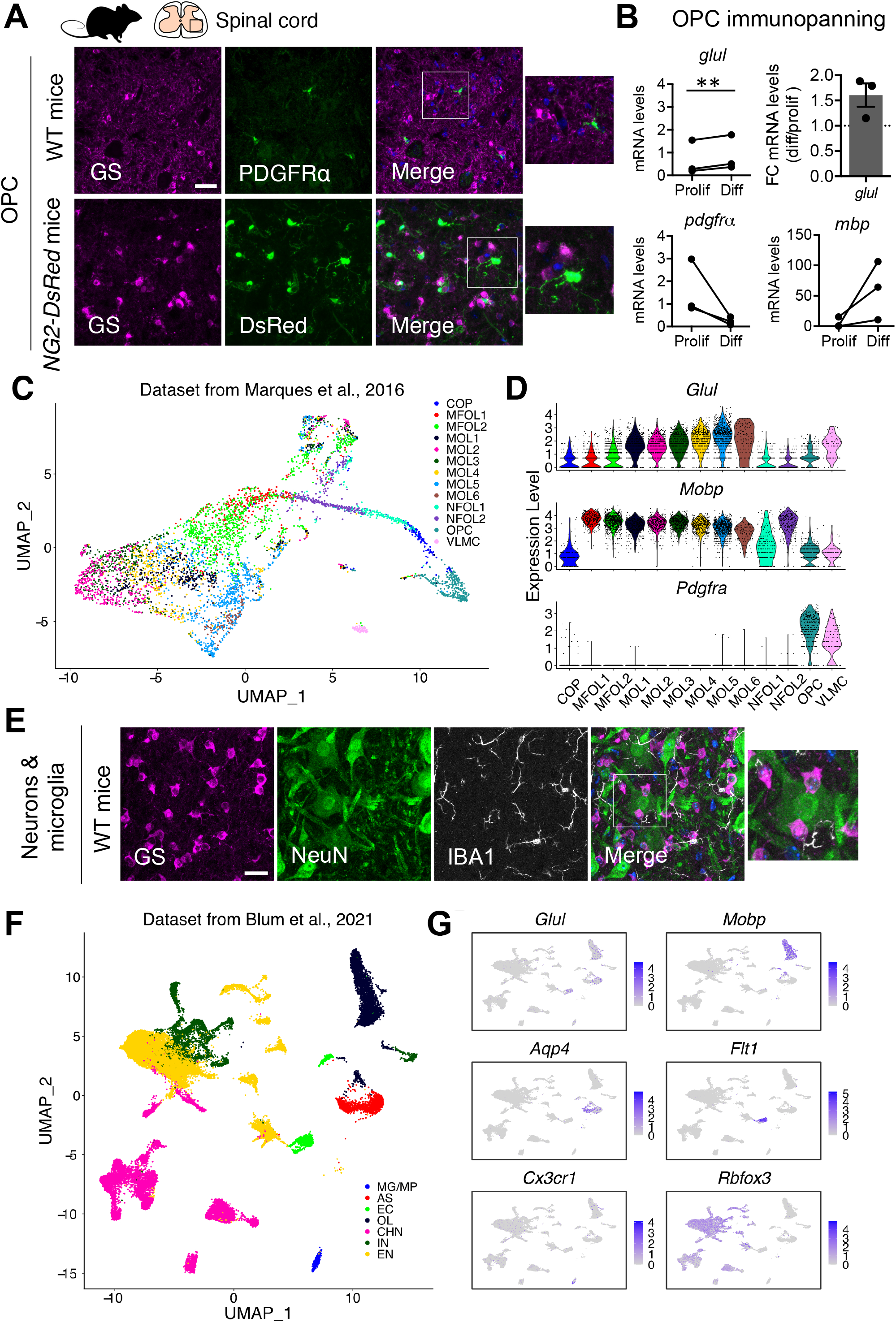
Spinal cord GS expression is macroglial-specific. **A,** Immunofluorescence costaining of GS (magenta) and OPC marker PDGFRα (green) in WT mice and in *NG2- DsRed* OPC reporter mice. **B**, mRNA levels and fold change (FC) of *Glul* (GS-encoding gene) between the proliferation (prolif, OPC) and differentiation (diff, OL) conditions. *Pdgfra* and *Mbp* were used as positive control markers for proliferation (OPC) and differentiation (OL) conditions. Data are expressed as mean ± SEM (n = 3 technical replicates). Paired t- test (t = 11.34, df =2), p < 0.001.**C,** Uniform Manifold Approximation and Projection (UMAP) plot depicting 5053 cells partitioned into 13 cell type and subtype clusters. VLMC: vascular and leptomeningeal cells, OPC: Oligodendrocyte progenitor cells, COP: Oligodendrocyte precursor, NFOL: newly formed OL, MFOL: Myelin-forming OL, MOL: Mature OL. **D**, Expression levels of *Glul*, *Mobp* and *Pdgfra*, in cell clusters identified in **C**. **E**, Immunofluorescence co-staining of GS (magenta), NeuN (neurons, green) and IBA1 (microglia, white). GS signal is not detected in neurons or microglia. **F,** UMAP depicting 43890 cells partitioned in 7 cell type clusters. MG/MP: microglia/macrophage, AS: astrocytes, EC: endothelial cells, CHN: cholinergic neurons, IN: inhibitory neurons, EN: excitatory neurons. **G**, Average expression of *Glul* and cell-type specific markers *Mobp* (OL), *Aqp4* (AS), *Flt1* (EC), *Cx3cr1* (MG) and *Rfbox3* (all neurons). Scale bars: 20μm (**A, D, E**).

Of note, the pattern of GS expression in OL is conserved between mouse and human CNS. Analyzing a published snRNA-seq data sets from human leukocortical tissues (Schirmer et al., 2019), we found that *GLUL* is expressed both in astrocytes and mature OL (**Supp. Figure 2A**). This was confirmed at the protein level using immunofluorescence co-stainings for GS and NOGO-A^+^, an established OL marker, in the ventral horn of human spinal cord sections (**Supp. Figure 2B**). Together, these results show that GS expression is macroglial-specific in the ventral spinal cord and that in oligodendroglia, it is expressed in mature OL rather than OPC.

### Topographical distribution of GS^+^ OL in the spinal cord

We next examined the distribution of GS^+^ OL in the mouse spinal cord. As shown in **Figure 3A, B**, GS^+^ OL are present along the spinal cord rostro-caudal axis at lumbar and cervical spinal cord levels. To determine if the density of GS^+^ OL was enriched in specific spinal cord areas, we quantified GS immunoreactivity in the dorsal gray matter (DGM), the ventral gray matter (VGM), the dorsal white matter (DWM) and ventral white matter (VWM) (**Figure 3C**). We measured GS maximum fluorescence intensity of GS^+^ signal in the different compartments, as GS^+^ OL cell bodies express GS at higher levels than the astrocytic GS (background) (see (**Figure 1A**)). GS fluorescence signal was more intense in GM versus WM areas and in VGM as compared to DGM, relative to the GFP signal in *MOBP-eGFP* transgenic OL line, in which the reporter is uniformly expressed throughout the spinal cord (**Figure 3D, E**).

**Figure 3.**
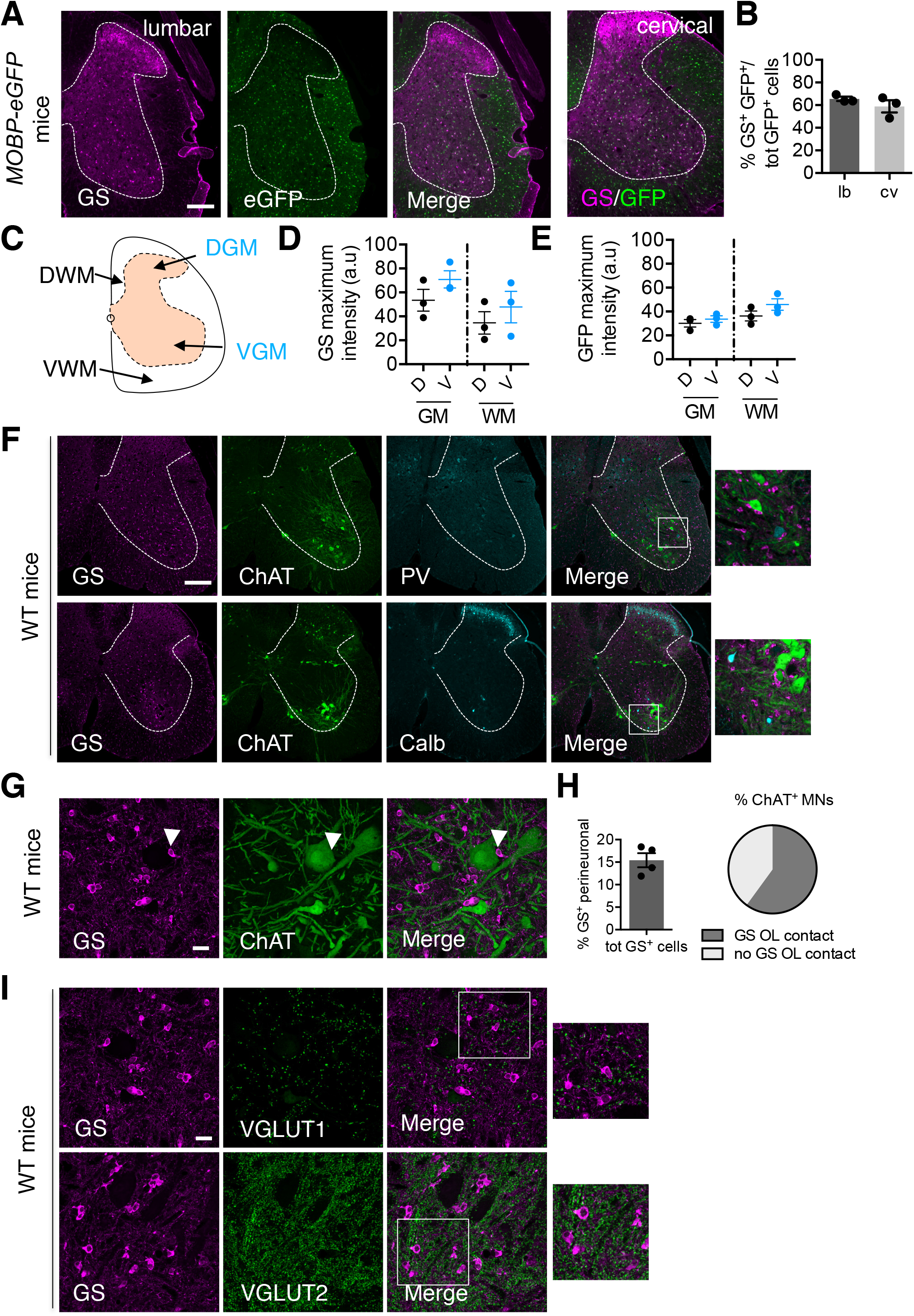
Distribution of GS^+^ OL in the mouse spinal cord. **A**, Low magnification epifluorescence images of spinal cord sections from lumbar (lb, left) and cervical (cv, right) levels from *MOBP-eGFP* mice stained with GS (magenta). **B**, Quantification of the % of GS^+^OL in the lb and cv spinal cord levels. **C-E,** Regions of interest in the dorsal (D) and ventral (V) grey (G) and white (W) matter (M) that were used to quantify GS (**D**) and GFP (**E**) maximum signal intensity. **F**, Confocal images of WT mice spinal cord sections coimmunostained with GS (magenta), the MN marker ChAT (green) and interneuron markers parvalbumin (PV) and calbindin (Calb) (cyan). **G**, High magnification confocal images of GS (magenta) and ChAT (green) showing GS^+^ OL perineuronal localization (white arrowhead). **H,** Quantification of the % of GS^+^ OL with a perineuronal localization and of the % of ChAT^+^ MN with or without contact with GS^+^ OL. **I**, Confocal images of fluorescent co-staining of GS (magenta) and glutamatergic terminals (VGLUT1 and VGLUT2, green). There is no specific organization of GS^+^ OL in relation to glutamatergic synaptic terminals in the ventral horn. Data are expressed as mean ± SEM (n = 3 - 4 mice). Scale bars: 200μm (**A**, **F**), 20μm (**G, I**).

Interestingly, GM GS^+^ OL often displayed perineuronal localization. Yet, this does not seem to be neuron subtype-specific as GS^+^ OL were found around ChAT^+^ motor neurons (MN), Parvalbumin^+^ (PV) and Calbindin^+^ (Calb) interneurons (**Figure 3F**). We found that more than half of the total MN cell bodies are contacted by GS^+^ OL, which represent about 15% of all GS^+^ OL (**Figure 3G, H**). As GS metabolizes Glu, we next determined if GS^+^ OL would topographically cluster according to glutamatergic terminals that are abundant in the mouse spinal cord. However, using immunofluorescence co-stainings of GS and glutamatergic terminal markers VGLUT1 and VGLUT2, we found no co-clustering of GS^+^ OL with glutamatergic synapses (**Figure 3I**).

### OL-specific GS loss of function does not lead to prominent histological alterations

We next used a conditional knockout (*cKO*) approach to investigate the functional role of *Glul* in spinal cord OL. We used *CNP-cre* to selectively remove *Glul* in mature OL, by breeding *Glul ^fl/fl^ mice* with *CNP-cre* mice, as previously described (Schirmer et al., 2018), (**Figure 4A**). *CNP-cre ^+^: Glul ^fl/fl^* (hereafter called *GS cKO*) mice showed almost complete elimination of GS^+^ OL in the mouse ventral horn as compared to *CNP-cre*^-^ (controls) (**Figure 4B**). Despite this, the ratio of Olig2^+^/CC1^+^ cell numbers reflecting OL differentiation was not different between *GS cKO* and controls, suggesting that it was not impacted by the loss of GS expression in OL (**Figure 4C**).

**Figure 4.**
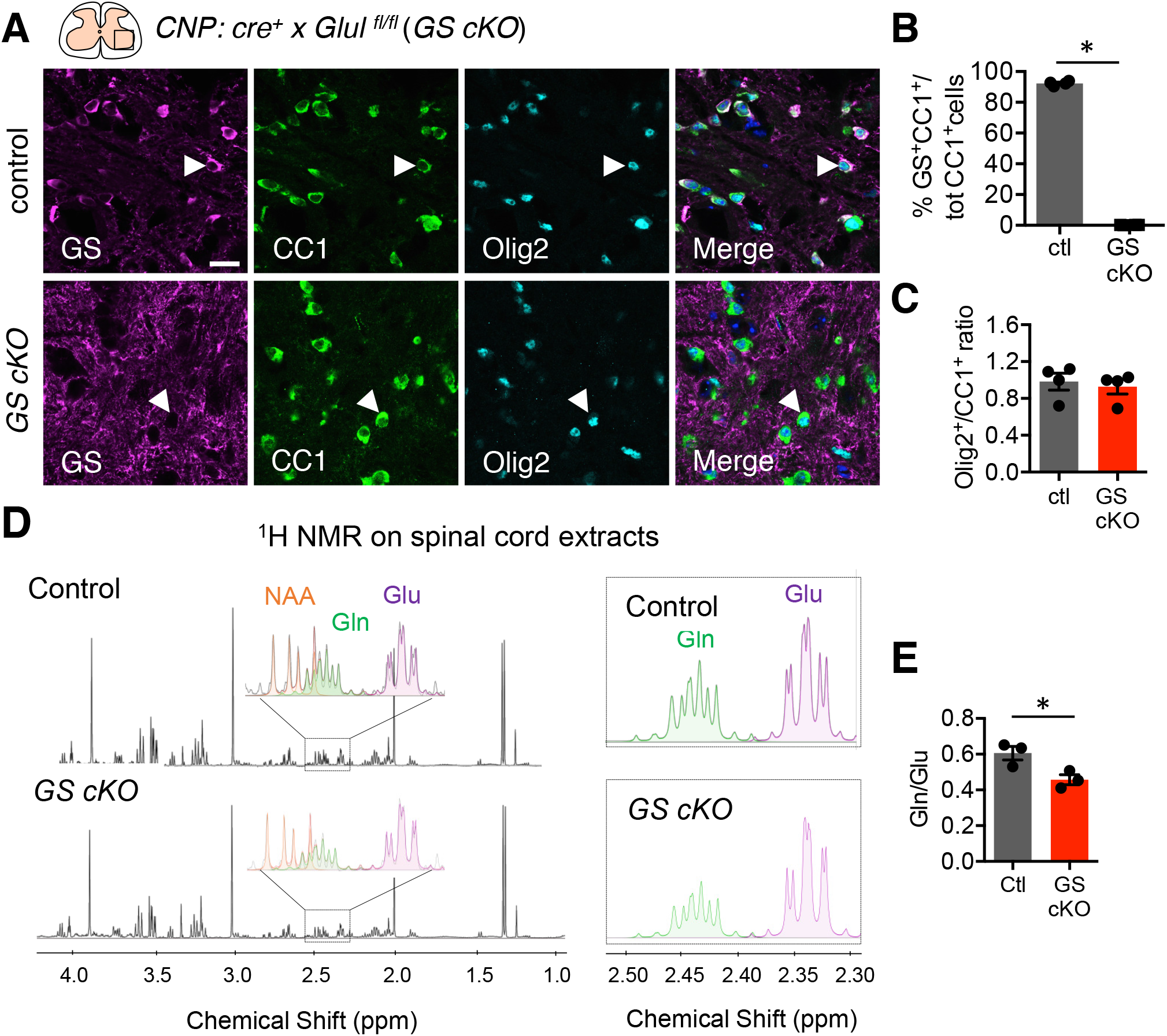
OL specific GS loss-of-function decreases glutamine levels in spinal cord extracts. **A**, Confocal images of spinal cord sections from *CNP-cre^-^: GS ^fl/fl^* (control) and *CNP-cre^+^: GS ^fl/fl^* (*GS cKO*) co-stained with GS (magenta) and OL markers CC1 (green) and Olig2 (cyan). **B**, Quantification of the % GS^+^CC1^+^/ tot CC1^+^ cells showing the drastic loss of GS expression in CC1^+^ OL. Data are expressed as mean ± SEM (n = 4 mice/group). Mann-Whitney test, p < 0.05. **C**, The Olig2^+^/CC1^+^ ratio, an index of OL differentiation is not different in 1 month-old (mo) *GS cKO* as compared to controls. **D**, Representative ^1^H NMR spectra showing relative abundance of metabolites from spinal cord extracts from 2 mo controls (top) and *GS cKO* (bottom) mice. NAA: N-acetyl-aspartate, Gln: glutamine, Glu: glutamate. **E**, Quantification of the Gln/Glu ratio showing that Gln levels are decreased in *GS cKO* mice. Data are expressed as mean ± SEM (n = 3 mice/group). Unpaired t-test (t = 3.159, df = 4), *p < 0.05.

In forebrain, we observed no change in GS expression in the cortex as GS is vastly expressed by astrocytes, which are not targeted with *CNP-cre*. However, in the pons, where GS is found in OL, a strong decrease in the number of bright GS^+^ cell bodies was observed (**Supp Figure 3A**). Consistent with histological data, GS protein levels, as detected by western blotting on ventral brain (pons and brainstem) samples, was decreased but not abolished since GS is still expressed by astrocytes (**Supp Figure 3B**). Furthermore, in the spinal cord, *GS cKO* efficiency was stable in time as similar results were obtained after 1 and 6 months of age (mo) (**Supp Figure 3C**).

GS metabolizes Glu into Gln. To determine whether GS loss in OL changed Glu and Gln levels, we used ^1-^H nuclear magnetic resonance (NMR)- based metabolomic analysis of spinal cord extracts from 2 months old *GS OL cKO* and control mice (**Figure 4D**). We found the Gln/Glu ratio was significantly decreased (**Figure 4E**). This result confirms that GS loss of function in OL has a functional role in metabolizing Glu into Gln.

We next investigated the effect of GS loss of function at the cellular level in the ventral spinal cord sensorimotor circuit. As shown **Figure 5A-C**, we found no difference in the numbers of ChAT^+^ MN (**Figure 5B**) or MN soma area (**Figure 5C**) either at 1 or 6 months. Ventral horn MN did not show signs of acute dysfunction as we did not observe enhanced neurofilament phosphorylation in neuronal cell bodies (SMI32 immunoreactivity), a marker of dysfunction (**Figure 5D**). Conversely, we did not detect a loss of axonal neurofilament phosphorylation by SMI312 immunoreactivity or myelin MBP expression in white matter tracts at 1 or 6 months (**Figure 5E-G**). Consistent with this result, we found no evidence of neuroinflammation in the ventral spinal cord (**Supp. Figure 3D**). Indeed, immunoreactivity levels of GFAP^+^ astrocytes and IBA1^+^ microglia (**Supp. Figure 3E, F**) were comparable in between *GS OL cKO* and control mice. Together, these results showed that the loss of GS expression in OL does not trigger detectable histological alterations as late as 6 months.

**Figure 5.**
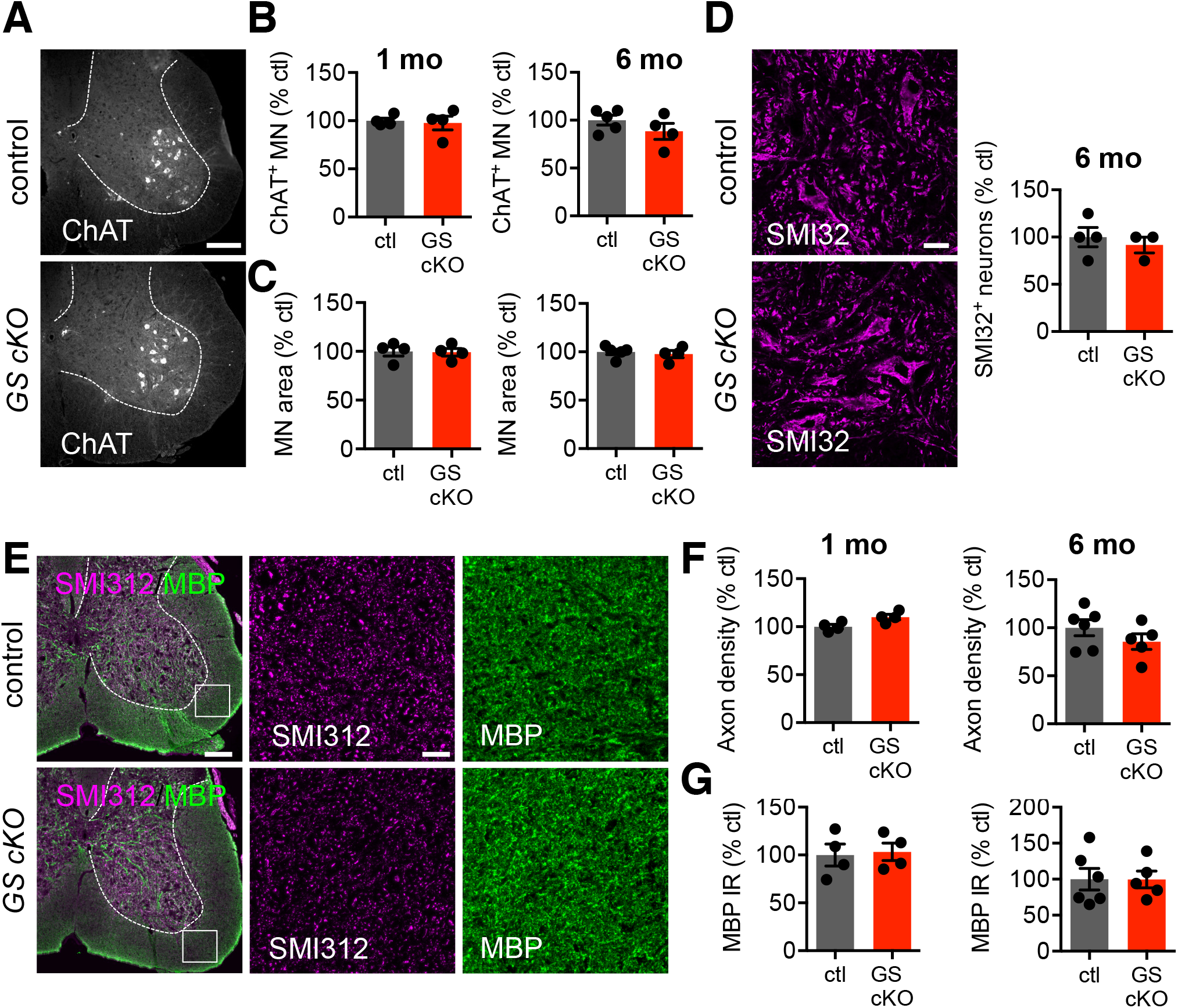
Loss of GS function in spinal OL does not lead to detectable histological changes from 1 to 6 months of age. **A**, Low magnification epifluorescence images of spinal cord sections from controls and *GS cKO* immunostained with the MN marker ChAT. **B**, **C**, Quantifications of the number (**B**) and the area (**C**) of ChAT^+^ MN in 1 and 6 mo mice. **D,** Immunofluorescence detection of the neuronal marker SMI32. No difference in SMI32^+^ neurons in 6mo *GS cKO* mice as compared with controls. **E,** Images of spinal cord sections from controls and *GS cKO* immunostained with the pan axonal marker SMI312 (magenta) and the myelin marker MBP (green). **F-G,** Quantification of axon density (**F**) and MBP immunoreactivity (**G**) in 1 and 6mo mice. Data are expressed as mean ± SEM (n = 4 - 6 mice/group). Scale bars: 20μm, 200μm.

### Transient decrease in muscle force in *GS OL cKO* mice

We investigated motor behavior in *GS cKO* mice and littermate controls from 2 to 6 months of age (**Figure 6A-C**). We first used the grip strength test to measure peak force in *GS cKO* and control mice at different time points (2, 4 and 6 mo) (**Figure 6A**). Male and female performances per genotype were not different, so both genders were pooled for each time points. Surprisingly, peak force was significantly decreased at 2 and 4 but this difference was not maintained at 6 mo. To determine whether this behavior was specific to peak force, we assess motor coordination and motor learning using the rotarod test (**Figure 6B**). We found no differences between the performances of *GS cKO* mice and controls at 2.5 and 4.5 mo. Finally, we used the open field test to assess general locomotor function (**Figure 6C**). We found that the total distance moved and number of rearings are not significantly different *GS cKO* mice and controls, even at 6 months of age. Together, these results showed that GS loss-of-function in OL results in a transient and specific loss of peak force in young mice.

**Figure 6.**
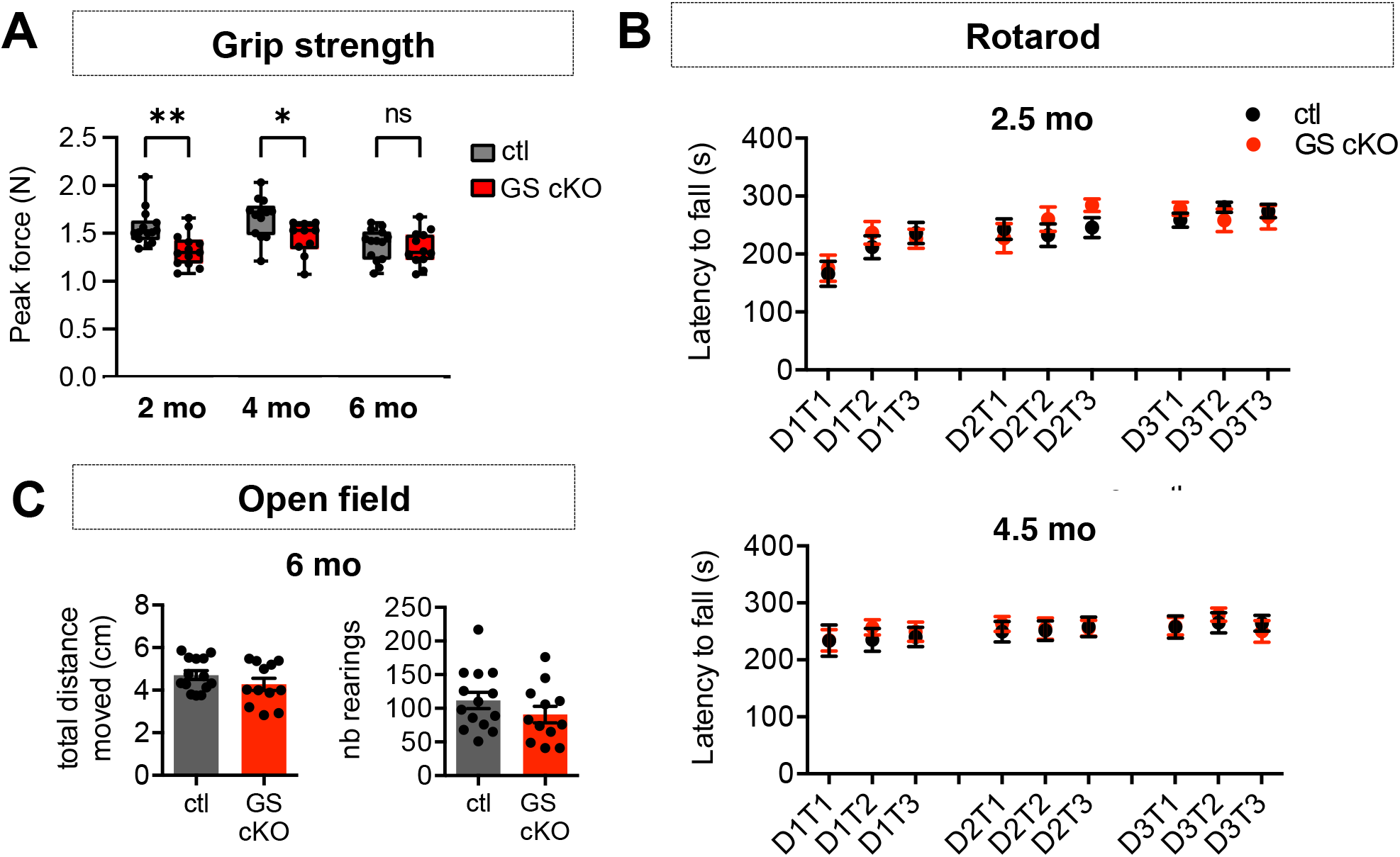
*GS OL cKO* mice show a transient loss of peak force but otherwise normal motor function. **A**, Peak force was measured using the grip strength test at 2, 4 and 6 months of age (mo). A transient decrease in peak force is observed at 2 and 4 months in *GS cKO* mice. In box-and-whisker plots, center lines indicate medians, box edges represent the interquartile range, and whiskers extend down to the 10th and up to the 90th percentiles of the distribution (n= 13-14 mice/group). Mixed Effect Model: genotype (F (1, 25) = 9,026, p < 0.01), time (F (2, 45) = 10,50, p < 0.001) and Sidak post hoc *p < 0.05, **p < 0.01. **B**, Rotarod performances of *GS cKO* and control mice at 2.5 mo (top) and 4.5 mo (bottom). **C**, At the open field test, *GS cKO* mice showed normal total distance moved and number of rearings, as compared to littermate controls at 6 mo.

## Discussion

This study is the first comprehensive characterization of GS-expressing OL in the spinal cord. Using a combination of immunofluorescence detection and cell type-specific markers or transgenic reporter mice, we found that GS is expressed in grey matter OL in both human and mouse spinal cord. Immunodetection of GS appears reliable for the following reasons: 1) the same results were obtained with two antibodies from two different suppliers, 2) it was validated using OL-specific *cre* recombination in *Glul ^fl/fl^* mice, where GS signal was completely abolished in OL but not in astrocytes (in two distinct OL-targeted cre lines: *CNP-cre* and *Olig2-cre* (data not shown)).

*GS cKO* mice show a significant decrease in peak force at 2 and 4 months of age. This was specific as motor coordination was not altered in *GS cKO* mice at 2.5 and 4.5 months. Because peak force is a specific of MN output as suggested by our previous work and others (Kelley et al., 2018; Muller et al., 2014), this result might suggest a specific impact of GS OL loss of function on MN. This timing coincides with the significant decrease of Gln levels in the spinal cord of *GS cKO* mice. Yet, this behavioral effect on peak force was not maintained at 6 months, suggesting that adaptive mechanisms operated by astrocytes might occur but take at least 4 months to be fully established.

We found that MN numbers, axon density and myelination were comparable in *GS cKO* and controls at both 1 and 6 mo. Although these histology-based approaches have been used in the past to evidence MN loss (Molofsky et al., 2014), they might not be sensitive enough to detect more subtle changes. It is possible that the observed functional effect is too transient to translate into histological modifications, which might occur over longer period of time or require more severe dysfunctions. In addition, it is possible that MN stress rather than death might be involved in *GS cKO* mice. Furthermore, it is likely that MN electrophysiological properties rather than numbers are altered in *GS cKO* mice. Unfortunately, it is experimentally very challenging to record MN activity using patch clamp in the adult mouse spinal cord. We and others performed electrophysiological recordings of spinal cord MN in mice as late as P15 (Kelley et al., 2018) but this timing is not compatible with GS expression onset in spinal cord OL, which occurs around P40.

We found that Gln/Glu ratio is decreased in spinal cord extracts from *GS cKO* mice, at 2 months of age using *ex vivo* ^1^H NMR. These results are consistent with previous findings obtained by HPLC in early postnatal glial *KO* mice (*human GFAP-Cre ^tg/-^/glul^fl/LacZ^*) (He et al., 2010). In another recent study, GS expression and function was characterized in midbrain OL using a distinct OL-targeting *cre* mouse line (*iMOG-cre*) (Xin et al., 2019). They evaluated Gln and Glu levels by colorimetric assay and found that both were significantly decreased in 2 mo *cKO* mice as compared to controls. These discrepancies could be explained because we used different methods for tissue measurement of metabolites and studied different CNS regions. Aside from this result, our findings are in accordance with this previous study. There was no detectable effect of GS loss of function on OL differentiation/myelination and these mice do not show drastic locomotor changes but subtle, cocaine-induced behavioral deficits.

Our findings add perspective on the functions of GS in macroglia. Similar to astrocytes, OL express glutamate transporters (Suárez-Pozos, Thomason, & Fuss, 2020). In culture, OL are able to uptake and rapidly metabolize excitatory amino acids (Reynolds & Herschkowitz, 1986; Tansey, Farooq, & Cammer, 1991). More recent work *in vivo*, showed that mature myelinating OL can provide metabolic to axons in the white matter through glutamate receptor-mediated signaling (Saab et al., 2016). In addition, grey matter OL with perineuronal localization may also provide metabolic support to neurons and regulate neuronal excitability in the mouse cortex (Battefeld, Klooster, & Kole, 2016). Together, these results suggest that, as in astrocytes, GS in OL could be involved in Glu-Gln cycle and metabolic support of neurons.

Not much is known about the function of grey matter OL in the adult mouse spinal cord, especially in pathological conditions. One elegant study found that in the superoxide dismutase SOD1^G93A^ mouse model of amyotrophic lateral sclerosis (ALS), which is characterized by the progressive loss of MN, adult-born grey matter OL accumulate in the ventral spinal cord (Gurney et al., 1994; Kang et al., 2013). Interestingly, in ALS, Glu-Gln cycle is altered and Glu-induced excitotoxicity is involved in neuronal death (Robberecht & Philips, 2013). Together with the present study providing evidence for a role of OL in Glu-Gln cycle, GS functions in grey matter OL might become more relevant in the context of ALS.

## Acknowledgements

We thank S. Fancy for providing *NG2-DsRed* mice. We also thank M. Shavali for technical help. We thank Drs. C. Escartin, C. Lobsiger and S. Boillee for inputs on the project, critical reading of the manuscript and feedback on figures. Authors were supported by Howard Hughes Medical Institute and the Wellcome Trust (D.H.R.), Paul G. Allen Foundation Distinguished Investigator Program (D.H.R.). L.S. was supported by research grants from the German Research Foundation (SCHI 1330/1-1), the Hertie Foundation (medMS MyLab, P1180016) and the National Multiple Sclerosis Society (FG-1607-25111 and FG-1902-33617). L.B.H. is currently supported by a Fondation pour la Recherche Medicale fellowship.

## Author contributions

L.B.H. conceived the project, designed and performed the experiments, wrote the manuscript and designed the figures. L.S. provided *CNP*-cre mouse breeders, human tissue, performed human tissue staining and imaging, helped with performing OPC immunopanning experiments and edited the manuscript. A.Z. performed bioinformatics analysis of published single cell/nucleus RNA-seq data setss and generated plots. B.T. performed ^1^H NMR experiments on spinal cord extracts. K.S. helped with behavioral testing and histological tissue processing. S.C. was in charge of mouse genotyping. W.H.L. provided the *Glul^fl/fl^* mice. M.M.C. analyzed ^1^H NMR data and edited the manuscript. D.H.R supervised the project and helped write the manuscript.

## Conflict of interests

Authors declare no conflict of interest.

## Notes

https://cells.ucsc.edu/

http://spinalcordatlas.org/

### Competing Interest Statement

The authors have declared no competing interest.

## References

Anlauf, E., & Derouiche, A. (2013). Glutamine synthetase as an astrocytic marker: its cell type and vesicle localization. Front Endocrinol (Lausanne), 4, 144. doi:10.3389/fendo.2013.00144

Battefeld, A., Klooster, J., & Kole, M. H. (2016). Myelinating satellite oligodendrocytes are integrated in a glial syncytium constraining neuronal high-frequency activity. Nat Commun, 7, 11298. doi:10.1038/ncomms11298

Bernstein, H. G., Bannier, J., Meyer-Lotz, G., Steiner, J., Keilhoff, G., Dobrowolny, H.,… Bogerts, B. (2014). Distribution of immunoreactive glutamine synthetase in the adult human and mouse brain. Qualitative and quantitative observations with special emphasis on extra-astroglial protein localization. J Chem Neuroanat, 61-62, 33–50. doi:10.1016/j.jchemneu.2014.07.003

Blum, J. A., Klemm, S., Shadrach, J. L., Guttenplan, K. A., Nakayama, L., Kathiria, A.,… Gitler, A. D. (2021). Single-cell transcriptomic analysis of the adult mouse spinal cord reveals molecular diversity of autonomic and skeletal motor neurons. Nat Neurosci, 10.1038/s41593-020-00795-0.

Cammer, W. (1990). Glutamine synthetase in the central nervous system is not confined to astrocytes. J Neuroimmunol, 26(2), 173–178. doi:10.1016/0165-5728(90)90088-5

Daikhin, Y., & Yudkoff, M. (2000). Compartmentation of brain glutamate metabolism in neurons and glia. J Nutr, 130(4S Suppl), 1026S–1031S. doi:10.1093/jn/130.4.1026S

Gong, S., Zheng, C., Doughty, M. L., Losos, K., Didkovsky, N., Schambra, U. B.,… Heintz, N. (2003). A gene expression atlas of the central nervous system based on bacterial artificial chromosomes. Nature, 425(6961), 917–925. doi:10.1038/nature02033

Gurney, M. E., Pu, H., Chiu, A. Y., Dal Canto, M. C., Polchow, C. Y., Alexander, D. D.,… et al. (1994). Motor neuron degeneration in mice that express a human Cu,Zn superoxide dismutase mutation. Science, 264(5166), 1772–1775.

Haberle, J., Gorg, B., Rutsch, F., Schmidt, E., Toutain, A., Benoist, J. F.,… Koch, H. G. (2005). Congenital glutamine deficiency with glutamine synthetase mutations. N Engl J Med, 353(18), 1926–1933. doi:10.1056/NEJMoa050456

Hafemeister, C., & Satija, R. (2019). Normalization and variance stabilization of single-cell RNA-seq data using regularized negative binomial regression. Genome Biology, 20(1), 296.

He, Y., Hakvoort, T. B., Vermeulen, J. L., Labruyere, W. T., De Waart, D. R., Van Der Hel, W. S.,… Lamers, W. H. (2010). Glutamine synthetase deficiency in murine astrocytes results in neonatal death. Glia, 58(6), 741–754. doi:10.1002/glia.20960

Jayakumar, A. R., & Norenberg, M. D. (2016). Glutamine Synthetase: Role in Neurological Disorders. Adv Neurobiol, 13, 327–350. doi:10.1007/978-3-319-45096-4_13

Kang, S. H., Li, Y., Fukaya, M., Lorenzini, I., Cleveland, D. W., Ostrow, L. W.,… Bergles, D. E. (2013). Degeneration and impaired regeneration of gray matter oligodendrocytes in amyotrophic lateral sclerosis. Nat Neurosci, 16(5), 571–579. doi:10.1038/nn.3357

Kelley, K. W., Ben Haim, L., Schirmer, L., Tyzack, G. E., Tolman, M., Miller, J. G.,… Rowitch, D. H. (2018). Kir4.1-Dependent Astrocyte-Fast Motor Neuron Interactions Are Required for Peak Strength. Neuron, 98(2), 306–319 e307. doi:10.1016/j.neuron.2018.03.010

Lappe-Siefke, C., Goebbels, S., Gravel, M., Nicksch, E., Lee, J., Braun, P. E.,… Nave, K. A. (2003). Disruption of Cnp1 uncouples oligodendroglial functions in axonal support and myelination. Nat Genet, 33(3), 366–374. doi:10.1038/ng1095

Liang, S. L., Carlson, G. C., & Coulter, D. A. (2006). Dynamic regulation of synaptic GABA release by the glutamate-glutamine cycle in hippocampal area CA1. J Neurosci, 26(33), 8537–8548. doi:10.1523/JNEUROSCI.0329-06.2006

Marques, S., Zeisel, A., Codeluppi, S., van Bruggen, D., Mendanha Falcao, A., Xiao, L.,… Castelo-Branco, G. (2016). Oligodendrocyte heterogeneity in the mouse juvenile and adult central nervous system. Science, 352(6291), 1326–1329. doi:10.1126/science.aaf6463

Mates, J. M., Campos-Sandoval, J. A., Santos-Jimenez, J. L., & Marquez, J. (2019). Dysregulation of glutaminase and glutamine synthetase in cancer. Cancer Lett, 467, 29–39. doi:10.1016/j.canlet.2019.09.011

Molofsky, A. V., Kelley, K. W., Tsai, H. H., Redmond, S. A., Chang, S. M., Madireddy, L.,… Rowitch, D. H. (2014). Astrocyte-encoded positional cues maintain sensorimotor circuit integrity. Nature, 509(7499), 189–194. doi:10.1038/nature13161

Muller, D., Cherukuri, P., Henningfeld, K., Poh, C. H., Wittler, L., Grote, P.,… Marquardt, T. (2014). Dlk1 promotes a fast motor neuron biophysical signature required for peak force execution. Science, 343(6176), 1264–1266. doi:10.1126/science.1246448

Norenberg, M. D., & Martinez-Hernandez, A. (1979). Fine structural localization of glutamine synthetase in astrocytes of rat brain. Brain Res, 161(2), 303–310. doi:10.1016/0006-8993(79)90071-4

Papageorgiou, I. E., Valous, N. A., Lahrmann, B., Janova, H., Klaft, Z. J., Koch, A.,… Kann, O. (2018). Astrocytic glutamine synthetase is expressed in the neuronal somatic layers and down-regulated proportionally to neuronal loss in the human epileptic hippocampus. Glia, 66(5), 920–933. doi:10.1002/glia.23292

Reynolds, R., & Herschkowitz, N. (1986). Selective uptake of neuroactive amino acids by both oligodendrocytes and astrocytes in primary dissociated culture: a possible role for oligodendrocytes in neurotransmitter metabolism. Brain research, 371(2), 253–266.

Robberecht, W., & Philips, T. (2013). The changing scene of amyotrophic lateral sclerosis. Nat Rev Neurosci, 14(4), 248–264. doi:10.1038/nrn3430

Rose, C. F., Verkhratsky, A., & Parpura, V. (2013). Astrocyte glutamine synthetase: pivotal in health and disease. Biochem Soc Trans, 41(6), 1518–1524. doi:10.1042/BST20130237

Saab, A. S., Tzvetavona, I. D., Trevisiol, A., Baltan, S., Dibaj, P., Kusch, K.,… Nave, K. A. (2016). Oligodendroglial NMDA Receptors Regulate Glucose Import and Axonal Energy Metabolism. Neuron, 91(1), 119–132.

Schirmer, L., Mobius, W., Zhao, C., Cruz-Herranz, A., Ben Haim, L., Cordano, C.,… Rowitch, D. H. (2018). Oligodendrocyte-encoded Kir4.1 function is required for axonal integrity. Elife, 7. doi:10.7554/eLife.36428

Schirmer, L., Velmeshev, D., Holmqvist, S., Kaufmann, M., Werneburg, S., Jung, D.,… Rowitch, D. H. (2019). Neuronal vulnerability and multilineage diversity in multiple sclerosis. Nature, 573(7772), 75–82. doi:10.1038/s41586-019-1404-z

Stuart, T., Butler, A., Hoffman, P., Hafemeister, C., Papalexi, E., Mauck, W. M.,… Satija, R. (2019). Comprehensive Integration of Single-Cell Data. Cell, 177(7), 1888–1902.e1821.

Suárez-Pozos, E., Thomason, E. J., & Fuss, B. (2020). Glutamate Transporters: Expression and Function in Oligodendrocytes. Neurochemical research, 45(3), 551–560.

Takasaki, C., Yamasaki, M., Uchigashima, M., Konno, K., Yanagawa, Y., & Watanabe, M. (2010). Cytochemical and cytological properties of perineuronal oligodendrocytes in the mouse cortex. Eur J Neurosci, 32(8), 1326–1336. doi:10.1111/j.1460-9568.2010.07377.x

Tansey, F. A., Farooq, M., & Cammer, W. (1991). Glutamine synthetase in oligodendrocytes and astrocytes: new biochemical and immunocytochemical evidence. J Neurochem, 56(1), 266–272. doi:10.1111/j.1471-4159.1991.tb02591.x

Tyagi, R. K., Azrad, A., Degani, H., & Salomon, Y. (1996). Simultaneous extraction of cellular lipids and water-soluble metabolites: evaluation by NMR spectroscopy. Magnetic resonance in medicine, 35(2), 194–200.

Wishart, D. S., Feunang, Y. D., Marcu, A., Guo, A. C., Liang, K., Vázquez-Fresno, R.,… Scalbert, A. (2018). HMDB 4.0: the human metabolome database for 2018. Nucleic acids research, 46(D1), D608–D617.

Xin, W., Mironova, Y. A., Shen, H., Marino, R. A. M., Waisman, A., Lamers, W. H.,… Bonci, A. (2019). Oligodendrocytes Support Neuronal Glutamatergic Transmission via Expression of Glutamine Synthetase. Cell Rep, 27(8), 2262–2271 e2265. doi:10.1016/j.celrep.2019.04.094

Yuen, T. J., Silbereis, J. C., Griveau, A., Chang, S. M., Daneman, R., Fancy, S. P.,… Rowitch, D. H. (2014). Oligodendrocyte-encoded HIF function couples postnatal myelination and white matter angiogenesis. Cell, 158(2), 383–396. doi:10.1016/j.cell.2014.04.052

