## Supplementary figures and images for "Evidence for glutamine synthetase function in mouse spinal cord oligodendrocytes"

### Supplemental Figure 1

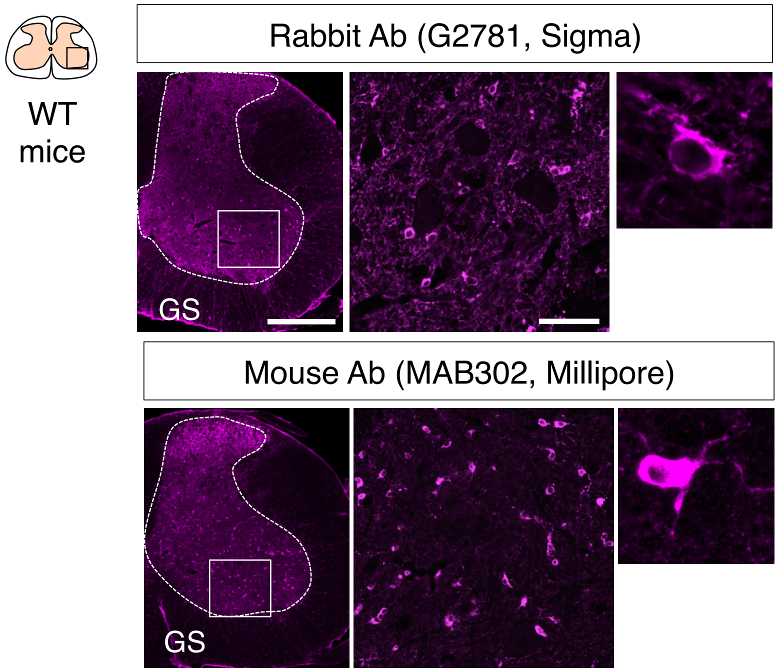

### Supplemental Figure 2

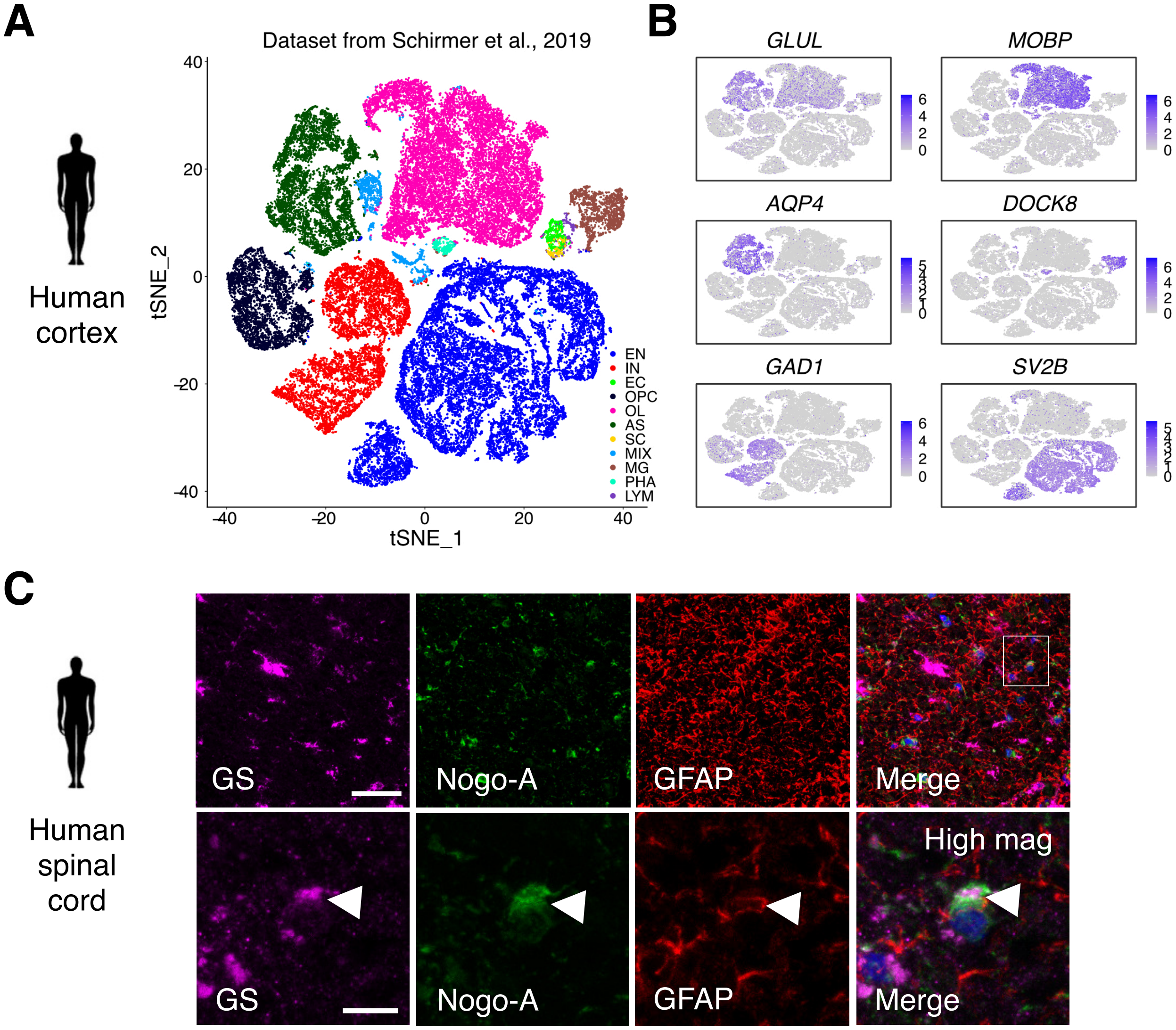

### Supplemental Figure 3

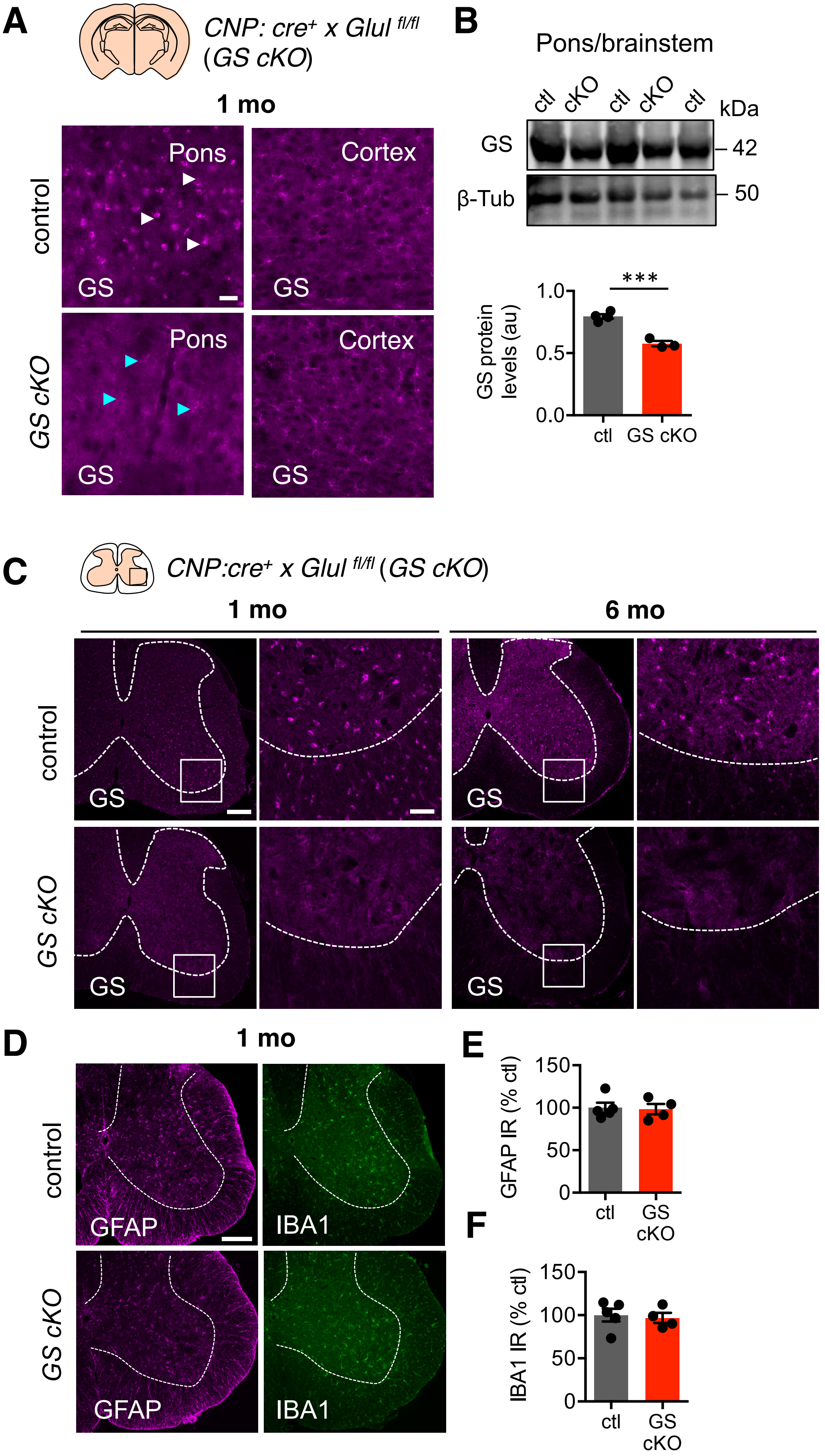
