## Supplemental Table 1 for "Evidence for glutamine synthetase function in mouse spinal cord oligodendrocytes"

| UK MS Tissue Bank # | Spinal cord tissue block # | RIN | Age | Sex | Postmortem interval (hours) |
|---|---|---|---|---|---|
| CO34 | SC6 | 6,7 | 88 | female | 20 |
| CO40 | SC4 | 7,1 | 86 | female | 29 |
| PDCO40 | SCB6 | 6,9 | 61 | female | 15 |

| Cause of death |
| --- |
| Pneumonia |
| Stroke |
| Ovarian cancer |
