## Supplemental Table 2 for "Evidence for glutamine synthetase function in mouse spinal cord oligodendrocytes"

| Present in matrix but not in metadata | Present in metadata but not in matrix |
|---|---|
| C1.1772099.081.D11 | C1.1772060.226.C07 |
| C1.1772099.081.G04 | C1.1772062.115.A10 |
| C1.1772099.081.B04 | C1.1772063.061.D10 |
| C1.1772099.081.F08 | C1.1772072.240.F10 |
| C1.1772099.071.A10 | C1.1772072.242.F03 |
| C1.1772099.081.H11 | C1.1772094.125.D10 |
| C1.1772099.081.H02 | C1.1772096.084.E05 |
| C1.1772099.081.G07 | C1.1772096.084.H03 |
| C1.1772099.081.F01 | C1.1772096.086.E08 |
| C1.1772099.071.B04 | C1.1772096.087.A03 |
| C1.1772099.071.B10 | C1.1772096.088.B11 |
| C1.1772099.071.B12 | C1.1772096.088.C03 |
| C1.1772099.071.D12 | C1.1772096.099.B12 |
| C1.1772099.071.B11 | C1.1772099.084.G10 |
| C1.1772099.071.C04 | C1.1772099.092.C12 |
| C1.1772099.071.A02 | C1.1772099.152.G03 |
