## Supplemental Figure legends for "Evidence for glutamine synthetase function in mouse spinal cord oligodendrocytes"

**Supplemental Figure 1. Antibodies for GS detection used in the study.** Low (left) and high magnification (right) images of GS immunofluorescence staining on mouse ventral spinal cord sections with two different antibodies. Scale bars: 200μm (low magnification, left) and 20μm (high magnification, right).

**Supplemental Figure 2. *GS-encoding gene* is expressed in mature OL in human samples. A**, UMAP plot depicting 48919 cells partitioned into 11 cell type clusters. SC: Stromal cells, MIX: mixture of glial cells, PHA: phagocytes, LYM: lymphocytes **B**, Expression levels of *GLUL*, and cell type specific marker genes *MOBP* (OL, oligodendrocytes), *AQP4* (AS, astrocytes), *DOCK8* (MG, microglia), *GAD1* (IN, inhibitory neurons), *SV2B* (EN, excitatory neurons). **C**, Confocal images of spinal cord sections from an adult human control (representative image selected from three control spinal cord cross-sections) immunostained with GS (magenta), the OL marker Nogo-A (green) and the astrocyte marker GFAP (red). Scale bars: 40μm (top), 10μm (high magnification, bottom).

**Supplemental Figure 3. Validation and stability of *GS cKO* in OL.** **A**, Epifluorescence images of brain sections from control and *GS cKO* mice showing the loss of GS expression in OL in the pons (white arrowheads) but not in the cortex, where GS^+^OL are sparse. Note the remaining expression of GS in pons astrocytes (cyan arrowheads). **B**, Western blot detection and quantification of 42-kDa GS protein on pons/brainstem brain samples from controls and *GS cKO* mice. β-III tubulin (β-Tub) was used as a housekeeping protein for protein level quantification. GS expression is decreased *GS cKO* mice as compared to controls (the remaining expression comes from astrocytes). Data are expressed as mean ± SEM (n = 3 - 4 mice/group). Unpaired t-test (t = 7.658, df = 5), ***p < 0.001. **C**, Low magnification epifluorescence images of GS staining in the ventral spinal cord of controls and *GS cKO* at 1 and 6 mo showing that *GS cKO* is stable in time. **D-F**, Low magnification epifluorescence images of GFAP (magenta) and IBA1 (green) in controls and *GS cKO* mice showing no evidence of astrocyte reactivity (quantified in **E**) or microglial activation (quantified in **F**) in 1 mo mice. **B**, Scale bars: 20μm (**A**, **C** high magnification), 200μm (**C** low magnification, **D**).

**Supplemental Table 1. Characteristics of human control spinal cord samples.**

**Supplemental Table 2. List of mismatched cells from the Marques et al., 2016 data set.**
